# A unified neurocomputational bilateral model of spoken language production in healthy participants and recovery in post-stroke aphasia

**DOI:** 10.1101/2020.02.21.959239

**Authors:** Ya-Ning Chang, Matthew A. Lambon Ralph

**Affiliations:** MRC Cognition and Brain Sciences Unit, University of Cambridge, UK

**Keywords:** stroke aphasia, language lateralisation, language recovery, bilateral language processing, neurocomputational modelling

## Abstract

Understanding the processes underlying normal, impaired and recovered language performance has been a long-standing goal for cognitive and clinical neuroscience. Many verbally-described hypotheses about language lateralisation and recovery have been generated. However, they have not been considered within a single, unified and implemented computational framework, and the literatures on healthy participants and patients are largely separated. These investigations also span different types of data, including behavioural results and fMRI brain activations, which augment the challenge for any unified theory. Consequently, many key issues, apparent contradictions and puzzles remain to be solved. We developed a neurocomputational, bilateral pathway model of spoken language production, designed to provide a unified framework to simulate different types of data from healthy participants and aphasic patients. The model encapsulates key computational principles (differential computational capacity, emergent division of labour across pathways, experience-dependent plasticity-related recovery) and provides an explanation for the bilateral yet asymmetric lateralisation of language in healthy participants, chronic aphasia after left rather than right hemisphere lesions, and the basis of partial recovery in patients. The model provides a formal basis for understanding the relationship between behavioural performance and brain activation. The unified model is consistent with the degeneracy and variable neuro-displacement theories of language recovery, and adds computational insights to these hypotheses regarding the neural machinery underlying language processing and plasticity-related recovery following damage.

**Significance Statement:** Studies of healthy and impaired language have generated many verbally-described hypotheses. Whilst these verbal descriptions have advanced our understanding of language processing, some explanations are mutually incompatible and it is unclear how they work mechanistically. We constructed a neurocomputational bilateral model of spoken language production to simulate a range of phenomena in healthy participants and patients with aphasia simultaneously, including language lateralisation, impaired performance after left but not right damage, and hemispheric involvement in plasticity-dependent recovery. The model demonstrates how seemly contradictory findings can be simulated within a single framework. To our knowledge, this provides the first coherent mechanistic account of language lateralisation and recovery from post-stroke aphasia.

## Introduction

Language is a key human ability and when impaired (e.g., after stroke or neurodegeneration), patients are left with significant disability. Aphasia (acquired language impairments that follow from brain injury, affecting comprehension, production, reading and writing) is common (1). Studies of healthy and impaired language have a long history, and these vibrant literatures have generated many verbally described hypotheses. The long-standing literature on aphasia dates back to seminal 19th century studies (2–4). While these verbally-described hypotheses advanced our understanding of language processing both theoretically and clinically, it is not clear how they work mechanistically and they can be mutually incompatible. For instance, some notions propose good aphasia recovery only results from language returning to the left hemisphere (5–9) while others report that recovered language performance is positively correlated with activation in the right hemisphere (10–12). As a recent review (13) noted, the current situation is confusing because there are many individual findings, different types of data (e.g., patients’ language performance vs. fMRI activations) yet no unified mechanistic account. There is a pressing need to have an implemented neurocomputational model which can provide: (a) a unified framework in which findings from healthy participants and aphasic patients can be accounted for; (b) a computationally-instantiated framework to formalise and test verbally-described hypotheses; and (c) a framework that can bridge between different types of cognitive neuroscience data including language behaviour, lesion locations and task-related fMRI. This was the overarching aim of the current study. The puzzles and targets are set out briefly below.

### Lateralisation assumptions

The first issue concerns lateralisation in healthy and impaired language. The very strongly held view that language is a left hemisphere function primarily arises from the long-standing neuropsychology literature showing that chronic aphasia is associated with left hemisphere damage but not with right hemisphere damage (14–16). However, the patient data are more graded than often portrayed. Recent evidence has shown that right hemisphere lesions can generate language problems especially in the early phase and some mild remaining deficits can be measured in chronic cases (17). Several patient studies of semantic cognition (18–20) also show that bilateral damage is required to show more substantial deficits.

Additionally, functional neuroimaging in healthy participants shows that many language tasks such as repetition, picture naming, comprehension and production might be bilaterally supported (21–26). Although the activation patterns are often leftward asymmetric, the degree of asymmetry largely depends on the nature of the tasks with a subset showing stronger forms of asymmetric bias. For instance, propositional speech production is more left lateralised whereas nonpropositional speech (e.g., counting) generates bilateral activations (27–29). Of course, identifying activations associated with language does not necessarily imply that the regions are necessary for language functions (30). Thus, it is important to note that a number of transcranial magnetic stimulation (TMS) studies of semantics (31–34) and phonology (35) also indicate that left and right areas contribute to healthy language.

Thus, when considering both chronic aphasic patients and healthy participants, it appears difficult to reconcile the seemingly contradictory findings: how can the language network be strongly left lateralised in patients but be bilateral, albeit asymmetric, in healthy participants? We propose that these results could reflect the outcome of an intrinsically bilateral yet asymmetric language network. Indeed, it has been demonstrated that functional asymmetry could follow from hemispheric asymmetry in the healthy language system (36–42) and that, when the system is damaged (e.g., in patients with low-grade glioma), the degree of asymmetry can change through functional plasticity (43, 44). For instance, using a combination of fMRI and diffusion tensor imaging, Vernooij et al (42) demonstrated a significant correlation between functional hemispheric lateralisation and the relative asymmetry of the arcuate fasciculus in healthy participants (see also (41)). Thus, within the language network, the majority of healthy participants show leftward asymmetry of brain volumes and arcuate fasciculus (36, 40, 45), suggesting that more of the computational capacity is in the left than the right hemisphere. Such capacity imbalance might generate bilateral yet asymmetric lateralisation in fMRI activation and also greater likelihood of chronic impairment after left than right damage. The latter may reflect a combination of the premorbid division of labour for left over right in healthy language, and the potential for plasticity-related recovery. This was explored in past computational work by re-exposing the damaged model to its learning environment, generating plasticity-related recovery via “retuning” of the remaining computational capacity (46, 47). A straightforward hypothesis, from these earlier models, is that the potential for such recovery reflects the amount of computational capacity available. Thus, if the right hemisphere has insufficient capacity to learn all language functions by itself *(see SI Appendix, S1)* then, when the dominant left hemisphere is entirely damaged, language functions cannot be fully re-established by the right hemisphere alone – c.f. chronic aphasia. In contrast, if the weaker right hemisphere is lesioned then the dominant left hemisphere may have sufficient spare capacity to assimilate the extra work.

### The computational bases of language recovery

A recent review (13) considered two mechanisms: *degeneracy* and *variable neuro-displacement*. Although not mutually-exclusive, degeneracy (30) suggests that cognitive functions might arise from multiple, structurally-distinct neural networks resulting in a partially resilient system. Following damage, recovery of function could be achieved by upregulation of quiescent regions, alternative pathways or non-language regions that are not typically engaged in the healthy state [for a computationally-implemented example, see: (46)]. The second mechanism is variable neuro-displacement, a concept borrowed from automotive engineering (variable displacement: (48)). Given that the brain is metabolically expensive, it seems very likely that energy consumption needs to be balanced against performance demand. This can be achieved in engines by ‘displacing’ (downregulating or turning off) a subset of cylinders when full power is not required. Returning to the brain, it is well-established that higher neural activity is coupled with increased metabolic energy costs (cf. neurovascular coupling). If we assume that a cognitive function is supported by a dynamic distributed network then, when performance demand is not maximal, parts of this network could be “displaced” (downregulated) to save energy (31, 49). This displaced ‘spare capacity’ is used when performance demand is high but it could also be permanently upregulated, after partial damage to the network, to support recovered performance (returning to the engine analogy, if one cylinder’s function was compromised then the other cylinders’ output could be upregulated to compensate).

Previous computational studies (46, 47) of plasticity-related recovery have provided some support for these principles, and highlighted two types of experience-dependent learning, each depending on remaining capacity in the model. In single pathway models, re-learning can retune and activate the ‘perilesional’ units and weights. Secondly, if there are multiple routes that support the task, re-learning can also shift the division of labour between different pathways. The potential for recovery-related changes is determined by the capacity available in different pathways and their engagement in the task prior to damage. Both mechanistic hypotheses about language recovery need to be specified in more detail within an implemented computational model that can simulate healthy and impaired language, as well as generate the different measures used to assess recovery of function, such as language performance and fMRI activations.

### Theories of aphasia recovery

In the long-standing literature on language recovery, most hypotheses are verbally described or are verbal descriptions of observed phenomena (13). For example, upregulated activation in perilesional and contralesional areas has been associated with recovered performance in post-stroke aphasia (5, 50–55). Van Oers et al. (54), for instance, showed that recovery of picture naming was associated with activation in the remaining portion of the left inferior frontal gyrus (IFG) while recovery on more demanding tasks was associated with upregulated contralesional activation in the right IFG in addition to the left IFG. There is also parallel evidence from combined TMS-fMRI studies in healthy participants that inhibition of left hemisphere regions upregulates activation in the right homologous regions (31, 32).

Another notion is the right hemisphere hypothesis (RHH). Several neuroimaging studies have demonstrated that patients with left hemisphere damage recruit the right hemisphere during language tasks (8, 51, 56). These findings have been interpreted in terms of a right hemisphere juvenile language system, which can provide some function after significant left hemisphere damage but it is generally weaker and error prone. Despite being a commonly repeated hypothesis dating over a century, the computational mechanisms involved in the development of language and the shifts of function after damage remain unspecified and computationally unimplemented. More confusingly, the hypotheses and data in relation to the RHH are contradictory. Some notions suggest that aphasia recovery is supported by this right hemisphere system: language performance is correlated with activation in the right hemisphere (10–12) and, when aphasic patients have a second right hemisphere stroke, their language performance declines (2, 57). In contrast, the ‘regional hierarchy framework’ proposes that right hemisphere activation is maladaptive resulting from a release of transcallosal inhibition, and good recovery only results from language returning to the left (5–9). In a seminal study of very mildly aphasic patients with good recovery (51), left hemisphere activation for auditory comprehension greatly decreased a few days after stroke, was followed by increased bilateral activation with a significantly upregulated peak in the right hemisphere two weeks after stoke, and then the peak activation shifted back to the left hemisphere in the chronic phase. However, it remains unclear what mechanisms underlie the changes in brain activity and what the longitudinal patterns are for moderate and severe aphasia.

These contradictory RHHs have inspired neurostimulation interventions with opposite aims: either promoting right hemisphere engagement (58) or trying to suppress it in favour of left hemisphere involvement (59–62). Without a better understanding of underlying mechanisms and a formal implemented model, various foundational issues remain. These include: how a right hemisphere system can develop if it is suppressed by the left hemisphere; how the two systems might interact; whether the results of negative associations between right hemisphere activation and language are simply a reflection of severity, as mild aphasia is associated with small lesions which leaves more of the left hemisphere intact and able to be activated. Our working assumption is that there is an intrinsically bilateral, albeit asymmetrically-provisioned single functional network. That is, the left and right hemispheres both contribute to speech production with differential contributions arising from the effects of imbalanced capacity across the hemispheres. An implemented model would permit a proper investigation of how this division of labour might shift and under what conditions, after brain damage.

Additionally, the hypothesis of the maladaptive right hemisphere activation supposes that the two hemispheres attempt to inhibit each other through transcallosal inhibition (6, 7, 9). There are several puzzles about this hypothesis including (a) why the healthy brain might spend most of its lifetime preventing regions from working (a biologically expensive implementation) and (b) how the less dominant system can develop even semi-useful representations if being persistently suppressed. We also note that, to the best of our knowledge, outside of the motor system (63–65) there are no demonstrations of transcallosal inhibitory connectivity. Conversely, there is even some evidence of excitatory connectivity (66, 67). With an implemented bilateral language model, we can explore the effect of transcallosal connectivity on model behaviour, task performance and recovery.

### Multiple measures

The last issue concerns different types of data and measures. Classically, explorations of brain function relied on relating brain activations (8, 10–12, 53) or lesions (68–70) to patients’ performance. Functional neuroimaging now allows the healthy and damaged brain to be explored, *in vivo*. Thus we now have multiple measures to consider in parallel, including lesion location and size, behavioural language performance, activations and connectivity. To make progress, the field needs to understand the relationship between these measures. It is tempting to assume that activated regions must be contributing to patients’ performance but activation does not prove necessity (30). Furthermore, different types of analyses, such as multiple voxel pattern analysis (71), have started to be used to explore and predict recovered performance. For instance, Fischer-Baum et al. (72) reported that, in a stroke patient with a severe reading impairment, the orthographic activation patterns in the right fusiform gyrus were more similar to stimulus patterns than in the left fusiform gyrus. Thus, it is critical that computational models are designed to accommodate multiple measures within a single framework, to allow formal explorations of the relationship between brain activations and contributions to the observed behavioural performance.

To summarise, the primary aim of this study was to address four key issues by developing a unified, bilateral pathway model of spoken language production: (a) language lateralisation in healthy participants and post-stroke patients; (b) mechanistic accounts for language recovery; (c) dynamic shifts of activation in post-stroke aphasia and recovery with/without transcallosal connectivity; and (d) the relationship between multiple measures and recovered function. We directly compared the model to data derived from four important, exemplar studies of healthy individuals and post-stroke aphasia: Vernooij et al. (42), Gajardo-Vidal et al. (17), Saur et al. (51) and Fisher-Baum et al. (72), and used the model to make novel predictions for future exploration. Given the maelstrom of historical hypotheses, data, etc., we have provided a summary guide to the critical issues, alternative viewpoints, our working hypotheses and simulated effects in Table 1.

**Table 1.** Overview of critical issues, key issues, alternative viewpoints, working hypotheses and simulated effects/model explorations.

| **Critical Issues** | **Key Issues** | **Alternative Viewpoints** | **Working Hypotheses** | **Simulated Effects/Explorations** |
| --- | --- | --- | --- | --- |
| ***Lateralisation assumptions*** | How can the language network be strongly left lateralised in patients but be bilateral, albeit asymmetric, in healthy participants? | (a) Impaired language function after left but not right hemisphere damage (14-16).<br>(b) Bilateral and asymmetric brain activations in healthy individuals during language tasks (21-29). | Leftward hemispheric asymmetry generates bilateral yet asymmetric lateralisation in simulated BOLD and also greater likelihood of chronic impairment after left than right damage. | (a) Functional asymmetry followed computational capacity; (b) A function-structure pattern (42); (c) Impaired performance after left but not right damage to the model (17). |
| ***The computational bases of language recovery*** | A lack of an implemented model for the computational bases of language recovery. | (a) Degeneracy (13, 30).<br>(b) Variable neuro-displacement (13). | The mechanisms are not mutually exclusive and they can be utilised as a part of the recovery process. | Various analyses on model behaviour and explorations of the underpinning computations in both damage and undamaged conditions. |
| ***Theories of aphasia recovery*** | (a) What are the dynamic activation shifts in post-stroke aphasia and recovery?<br>(b) What is the effect of transcallosal connectivity on healthy and impaired function? | (a) Perilesional upregulated activation (5, 50-55).<br>(b) Right hemisphere activation (2, 8, 10-12, 51, 56-57).<br>(c) The ‘regional hierarchy framework’ (5-9). | Patterns of recovery are related to the differential capacity available in left and right hemisphere systems and lesion severity. | (a) Dynamic patterns of activation shifts during recovery (51); (b) Lesion severity as a determiner for recovered performance and brain activation patterns; (c) Different types of interconnectivity. |
| ***Multiple Measures*** | What is the relationship between multiple brain measures and recovered function in patients? | (a) Behavioural measures compared with fMRI activation (8, 10-12, 53).<br>(b) Potential applications of multiple voxel pattern analysis (71, 72) in the patient studies. | Different measures provide different types of information. | (a) Model accuracy better tracked by the RSA than unit activation (cf. BOLD); (b) A conceptually similar RSA pattern to data in (72). |

## Results

### Hemispheric asymmetry and language lateralisation

The bilateral model was implemented as a simple recurrent network, consisting of two parallel pathways trained to perform word repetition (see Methods). We investigated if the model could simulate language lateralisation that follows hemispheric asymmetry, similar to the correlation pattern between functional hemispheric asymmetry in parietotemporal regions and structural asymmetry in arcuate fasciculus during the spoken production task reported in Vernooij et al (42). Specifically, we varied the proportion of hidden units in the left versus the right pathways in the model (see Fig. 1A) to simulate the relative capacity of the two pathways (73) while the total number of hidden units remained unchanged. The number of units for the two consecutive hidden layers in each pathway was the same. After training, the model was also tested on nonword repetition (an assessment of generalisation to novel phonological forms).

**Fig. 1.**
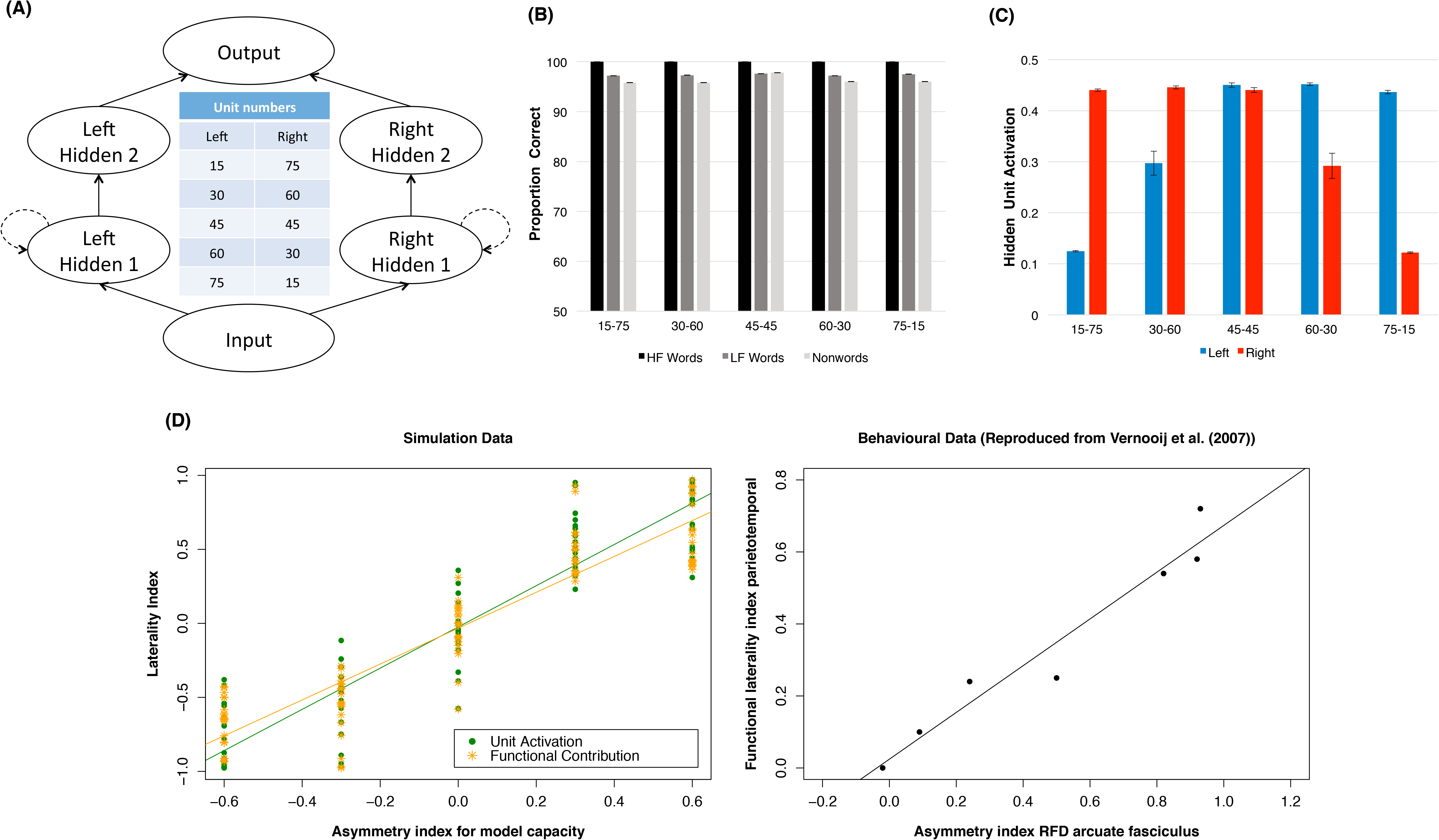
The model architecture, repetition performance, average hidden unit activation, and lateralisation patterns produced by the bilateral model with differential capacity in the left and right pathways. (A) The model with five different numbers of hidden units in the left and right pathways including 15-75, 30-60, 45-45, 60-30, and 75-15. The number of units in hidden 1 layer and hidden 2 layer was the same. The dashed lines indicate Elman connections (see Methods); (B) The repetition performance of the model on high frequency words, low frequency words and nonwords; (C) Hidden unit activation produced by the model across the hidden layers along the left and right pathways; (D) The lateralisation patterns based on functional contribution and output unit activation produced by the model and the behavioural data reproduced from Vernooij et al. (42). HF: high frequency; LF: low frequency. Error bar represents ±1 standard error; RFD: relative fibre density.

In imaging studies, a laterality index is commonly estimated using BOLD or cerebral blood flow (CBF) in the left and right homologue language areas (42, 74). In our model, two different measures were used to compute the degree of lateralisation: *functional contribution* and *output unit activation*. Functional contribution measured the relative contribution from the left or right pathway to output activation, as a proxy of effective connectivity analyses (47, 75). Output unit activation measured average unit activation at the output layer from either the left or right pathway, as a proxy of fMRI activation (76). A positive laterality score indicated that the model showed a left lateralised pattern; conversely, a negative score indicated a right lateralised pattern. An asymmetry index for computational capacity (number of left vs. right hidden units) was calculated in the same way as the laterality index. We also investigated average hidden unit activation in the left and right pathways across different conditions, during development and in recovery.

Results are summarised in Fig. 1. All models performed well on word repetition and generalised to nonwords (Fig. 1B). There was a clear lexicality effect with the highest accuracy for high frequency words followed by low frequency words and then nonwords. Importantly, the performance level achieved by the model with differential capacities in the left and right was very similar because the total number of units was the same. These observations were confirmed by a repeated ANOVA. There was a significant word type effect [HF vs. LF vs. nonwords: F(2, 278) = 33.8, *p* < .001], while both model type (*p* = 0.44) and its interaction with word type (*p* = 0.18) were not significant. This means that the model was able to exploit the computational capacity flexibly to learn the task and to generalise. In contrast, the underlying processing changed. Figure 1C shows that more hidden units along a pathway resulted in higher average hidden unit activation. Thus, the emergent functional division of labour in the model was not solely based on there being more units in the “dominant” processing pathway but they also resultantly worked harder on average. Critically, Figure 1D shows that the model with more processing units in the left pathway (i.e., a larger asymmetry index) produced a more left lateralised pattern. Laterality indices based on functional contribution and output unit activation were both positively correlated with asymmetry index for model capacity, Pearson’s *r* = 0.929, *p* < .001, and Pearson’s *r* = 0.917, *p* < .001, respectively. The results are consistent with the function-structure pattern reported in Vernooij et al. (42).

### Chronic aphasia after left but not right hemisphere stroke

We next investigated whether damage to the left hidden layer in the model would be more likely to result in permanently impaired language performance (chronic aphasia) compared to damage to the right. Gajardo-Vidal et al. (17) found that approximately half of patients 49% (151/307) with left hemisphere stroke showed impaired repetition performance, whereas the incidence for patients with right hemisphere stroke was about 5% (5/93). In the preceding section, we demonstrated that the model with more computational capacity in the left pathway produced a bilateral, left-asymmetric activation pattern similar to fMRI brain activations observed in most healthy individuals during language production. Thus, we opted to use a model with an asymmetrical structure where the computational capacity in the left was twice as large as that in the right (60 vs. 30 units). The 30 units in a hidden layer also met the minimum number of units required for a unilateral model to support the production task *(see SI Appendix, S1)*.

Fig. 2A shows both the developmental learning trajectory before lesioning (the intact model) and an example recovery profile of the model with a left or right moderate lesion. During development, the model learned high frequency words more accurately and quickly than low frequency words. Generalisation to nonwords was very good though lower than performance on words. Then, a moderate lesion was applied to the left or right hidden layer 1 in the model. A moderate lesion 50%[0.5] meant that 50% of the units were damaged and noise with the variance of 0.5 was added to the links connecting to and from the left hidden layer 1. After damage, the model was re-exposed to its learning environment for 100,000 word presentations to allow for a period of experience-dependent, plasticity-related recovery (based on a re-optimisation of the remaining processing units) (47). To mimic the loss of function and missing activation in the damaged brain regions immediately after stroke (51), a period of initial inefficient learning for the surviving units was implemented (i.e., their learning abilities were initially limited and then gradually regained, whereas for the units in the unaffected layers learning efficiency was normal). Inefficient learning was implemented by varying unit gain from 0 to 1 in steps of 0.1 over the early stage of retraining (the first 10,000 word presentations in the recovery phase). Note that the model behaved similarly without the implementation of a period of inefficient learning *(see SI Appendix, S2)*. Immediately after left damage, the performance of the model was at floor. Then, the model started to re-optimise the weight connections and re-learned the task. In the later stage of recovery, performance gradually increased up to an asymptote (i.e., partial function recovery as found in chronic aphasia). In contrast, the right damage only caused minor disruptions to the performance and it recovered rapidly (i.e., full function recovery akin to transient aphasia).

**Fig. 2.**
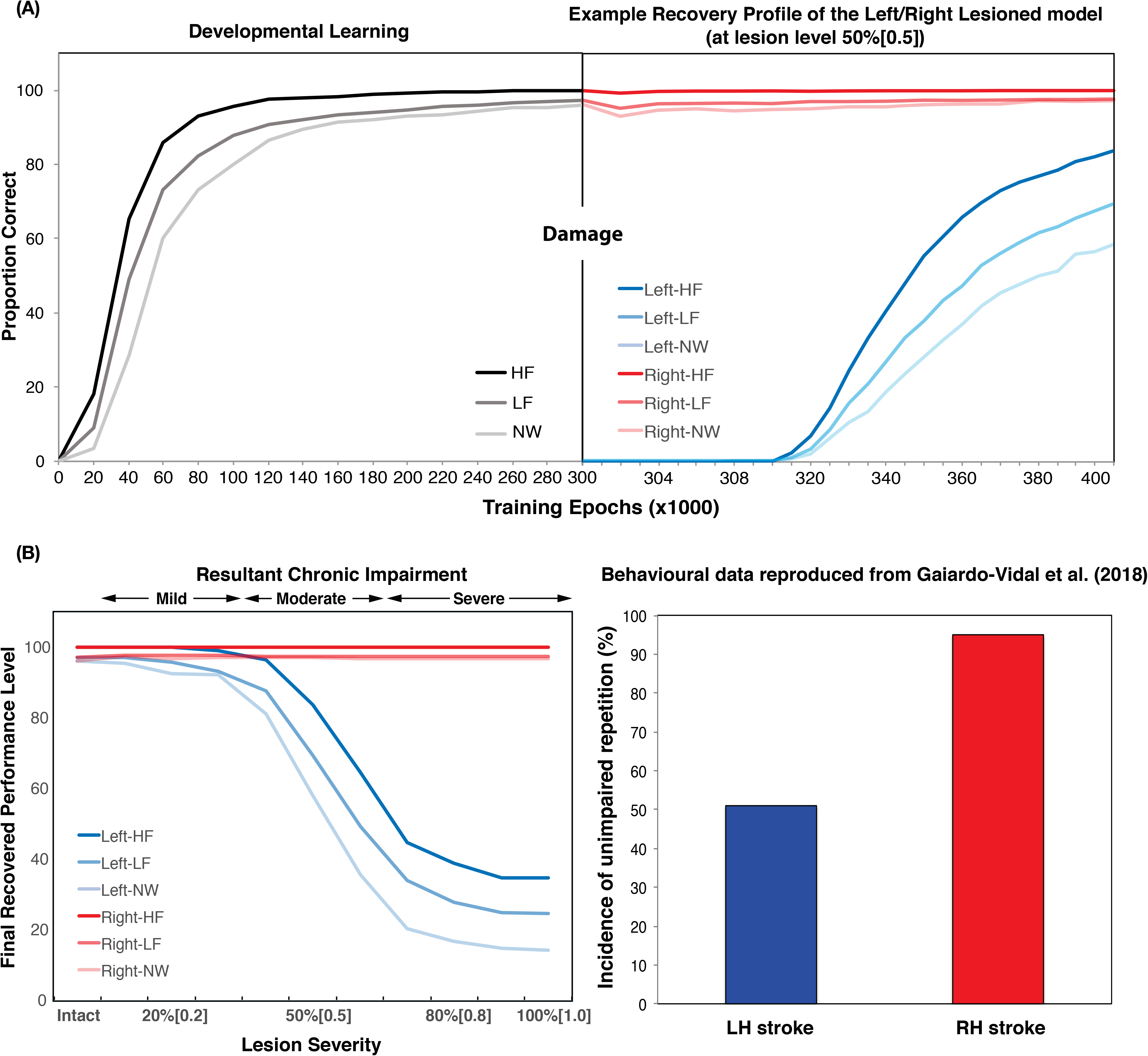
(A) The developmental learning trajectory of the model before damage, and the recovery profile after damage (Moderate lesion 50%[0.5]) to the left or right hidden layer 1, simulating a left or right hemisphere stroke and recovery. Note that the unequally spaced time scales for the re-learning period were made to clearly demonstrate the model’s re-learning in different periods; (B) The recovered performance of the left lesioned model and the right lesioned model as a function of lesion levels (a combination of unit damage and noise – see text for details). ‘Intact’ means the model without lesion. The behavioural data of unimpaired repetition performance were reproduced from Gajardo-Vidal et al. (17). HF: high frequency words; LF: low frequency words; NW: nonwords; LH: left hemisphere; RH: right hemisphere.

Obviously, patients have different lesion severities in the left or right hemisphere, leading to different recovery profiles. To capture this, different levels of damage were applied to the left or right hidden layer 1. Specifically, ten lesion levels were implemented by damaging hidden units from 10% to 100% with step increment of 10%, plus adding Gaussian noise with variance from 0.1 to 1 with step increment of 0.1 to the links that were connected to and from the target hidden layer. All re-training procedures were the same as described above. Fig. 2B shows the final recovered performance as a function of different levels of damage to the left or right hidden layer 1. For left lesions, the recovered performance varied with lesion levels. We divided the models into three lesion groups, 10%[0.1]-30%[0.3] for the mild group, 40%[0.4]-60%[0.6] for the moderate group, and 70%[0.7]-100%[1] for the severe group. The mild group showed the best recovered performance while the severe group was the worst with the moderate group in the middle. For the right lesions, the model generally recovered very well regardless of lesion levels. These results demonstrated that, following damage and recovery, performance of the left lesioned model was much more impaired than the right lesioned model. The simulation data were generally consistent with the patients’ studies reported in Gajardo-Vidal et al. (17), showing a stroke in the left hemisphere is more likely to lead profound, chronic language impairment (Fig. 2B), albeit the right lesion may have underestimated the mild level of aphasia that is sometimes observed (see Discussion).

### Dynamic activation shifts in post-stroke aphasia and recovery

An important aspect of this study was to investigate the relationship between simulated behavioural performance and underlying metrics of unit function (to mimic functional neuroimaging data). Three levels of left lesions (20%[0.2], 50%[0.5], and 80%[0.8]) were selected to simulate mild, moderate and severe aphasia. Additionally, the severe right lesion (80%[0.8]) was included to understand what compensated the effects of right damage. Four measures were used to reveal the mechanisms underlying recovery in the damaged model. First, as before, the damaged model’s accuracy on word and nonword repetition was used to simulate post-stroke aphasic patients’ behavioural performance. Second, we used output unit activation in the left and right pathways as a proxy of fMRI activation (76). Additionally, we investigated whether the model could produce similar activation patterns to that observed by Saur et al. (51) during three phases of language recovery. Specifically, we investigated whether a mildly lesioned model could produce: (a) from the acute phase to the subacute phase, an increase of the output unit activation in the undamaged left pathway and the right pathway with the highest increase in the right; (b) from the subacute phase to the chronic phase, a decrease of the output unit activation in the right pathway but the output unit activation in the left remained stable. Accordingly, the re-learning time in the model was divided into three recovery periods (acute, subacute, and chronic) approximating different stages of patient recovery, and the average output unit activations were computed. Third, we measured the perilesional and contralateral hidden unit activations to examine which undamaged units in the model were reformulated to support during recovery. Lastly, we conducted representation similarity analysis (RSA) comparing the activation similarity patterns in the hidden layers to the output similarity for the words. To our knowledge, there is only one stroke patient study that has utilised RSA (72). This study found that, when reading, a patient with a severe lesion to the left VWFA relied more on the right VWFA for orthographic processing, indexed by the RSA similarity scores, while the healthy participants generally relied more on the left VWFA than the right. As this investigation was a single case study of reading (not repetition), we report analogous data based on the closet settings to Fischer-Baum et al. (72) and investigated whether the reliance of the processing shifted from the left to right after severe damage to the left in the model.

In addition to these four measures, two additional measures were related to the model’s relearning: average weight strength and weight change. Both measures were helpful for understanding how the model re-learned the task during recovery and what the links were between recovery performance and re-learning processes *(see SI Appendix, S3 for details)*.

Fig. 3 summarises several key phenomena. We can first look at performance accuracy and output unit activation. For the left lesion, the recovered performance of the model aligned with lesion severity with the mild lesion model showing the best performance. Importantly, for the mildest lesion there was a transient pattern of output unit activation shifting from left to right at the early stage of recovery and then back to left at the later stage of recovery, similar to the finding observed in the mild aphasic patients (51). To test these observations, we compared the simulation data against Saur et al. (51) by computing the differential output unit activations between the acute and the subacute phases, and between the subacute and the chronic phases. As can be seen in Fig. 4A, the output unit activations significantly increased for both the left and right pathways, *t* = 8.11, *p* < .001, and, *t* = 8.56, *p* < .001, respectively, from the acute to the subacute phase. The increase of activation was numerically higher for the right than for the left pathway. The comparison of the subacute and chronic phases showed a significant decrease of activation in the right pathway, *t* = −3.55, *p* = 0.002, but not in the left pathway (*p* = 0.14). All the statistics were corrected for multiple comparisons.

**Fig. 3.**
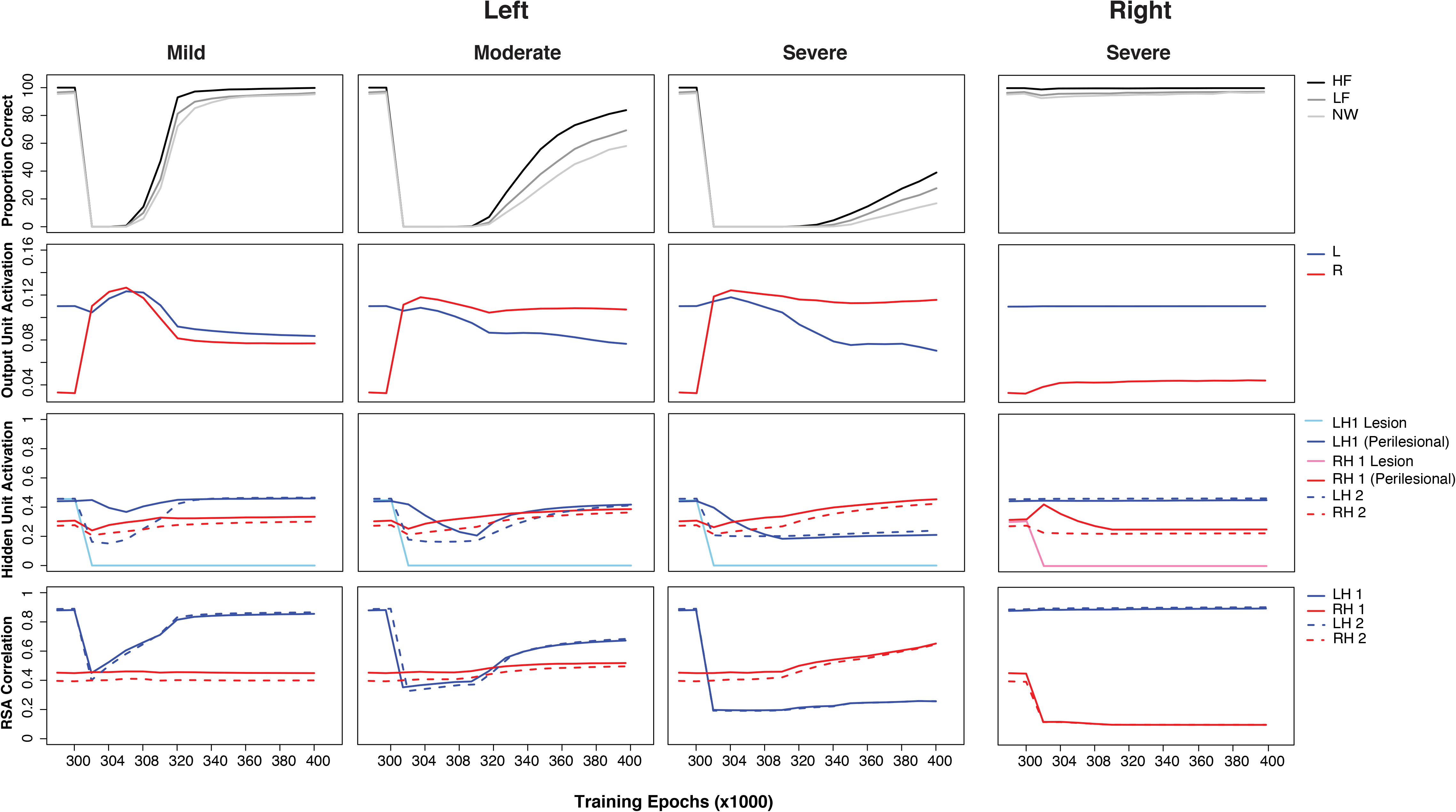
Simulation patterns of post-stroke aphasia and recovery: left mild (20%[0.2]), left moderate (50%[0.5]), left severe (80%[0.8]) and right severe (80%[0.8]) conditions. The lesion level was a combination of the proportion (%) of the units was damaged and the range of noise (bracket) added to the connections to and from the hidden layer. For each lesion condition, the first panel shows model performance; the second panel shows output unit activation generated from the left and right pathway of the model separately; the third panel shows hidden unit activation for the left and right hidden layers 1 and 2. The activations for lesioned and perilesional units are plotted separately; the last panel shows the RSA scores obtained in the left or right hidden layers 1 and 2 in the model. HF: high frequency words; LF: low frequency words; NW: nonwords; L: left; R: right; LH: left hidden layer; RH: right hidden layer.

**Fig. 4.**
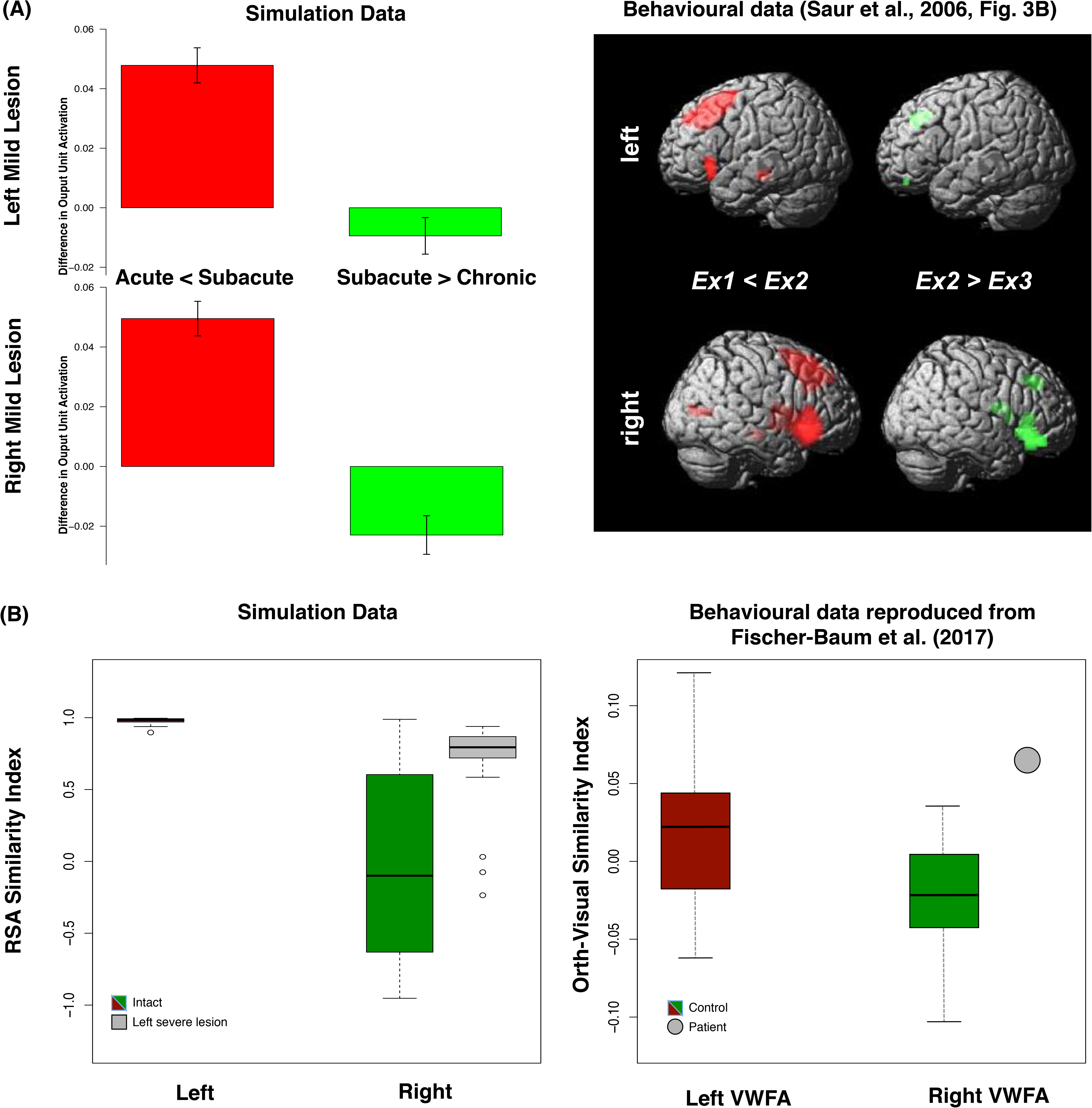
(A) The increment and decrement of the activations in the model with a left mild lesion. Direct comparisons of output unit activations between the acute (300K-301K) and subacute (301K-305K) phases and between the subacute and chronic (305K-340K) phases. The fMRI data of upregulation and downregulation in the language network from Saur et al. (51). Ex1: Acute; Ex2: Subacute; Ex3: Chronic. (B) RSA similarity indices of the left hidden layer 1 for both the intact model and the model with a severe left lesion. The orth-visual similarity indices for both the controls and the patient in the left VWFA and the right VWFA from Fischer-Baum et al. (72). VWFA: Visual word form area.

As for the moderate and severe lesion, the models in Fig. 3 showed right lateralised activation patterns, and the recovered performance was worse than that in the mild lesion. In contrast, even after a severe right lesion, accuracy was only slightly disrupted but quickly recovered, and the output activation pattern during recovery largely remained unchanged with a small rise in right output unit activation.

We investigated how undamaged perilesional and contralesional units supported recovery. The results showed that, for both mild and moderate lesions, the LH1 perilesional activation initially decreased following damage but then gradually increased during re-learning, reflecting a re-optimisation process. A similar but larger initial decrement followed by a slower increment pattern was observed for LH2 hidden unit activation. For a severe lesion, both the LH1 and LH2 hidden unit activation decreased following damage but did not rise again, presumably because there were insufficient processing units available in the LH1 layer for the model to re-optimise. This pattern was also observed for the right severe lesion comparison, where both the RH1 perilesional activation and RH2 hidden unit activation gradually decreased and remained in a low activity level. Turning to contralateral activation, for all severities of the left lesion, the contralateral hidden unit activations at RH1 and RH2 increased very quickly following damage. The degree of increment was varied and depended on lesion severity, with the largest increment for the severe condition. By contrast, for the right severe condition, there was no clear increment of the contralateral hidden unit activations at LH1 and LH2.

For the correct interpretation of the relationship between patient behavioural performance and underlying activation, it may be important to note that there were differential associations between model accuracy and the various unit metrics. Fig. 3 shows that the RSA measure closely shadowed the changing model accuracy, quite unlike simple unit activation (a proxy to BOLD levels) which show a complex nonlinear relationship. Taking the left moderate lesion as an example, even when the right output unit activation was building up quickly during the initial recovery period, change in model performance was minimal. Subsequently, long after the point when the right output unit activation reached a relatively stable level, there was a much larger and gradual increase in model accuracy. By contrast, the change in the RSA pattern was closely aligned with model performance.

Interestingly, although the right output unit activation was higher than the left output unit activation throughout recovery, the RSA results showed the left unit correlation was initially lower than the right unit correlation but returned to a higher level later in recovery. To examine, formally, the relationships between model performance with output unit activation and the RSA measure, we conducted correlation analyses. Model performance was correlated with output unit activation and the RSA scores at hidden layers 1 and 2 separately. Correlation analyses were conducted across the developmental learning period in the intact model and the re-learning period in the lesioned model. Results are summarised in Table 2. The correlations between output unit activation and model performance were mostly negative in particular for the lesioned conditions, except for the positive correlations for the left output unit activation in the intact condition and for the right output unit activation in the left severe lesion condition. When considering all intact and lesion conditions, the pattern of change in correlation for output unit activation was difficult to interpret. By contrast, the correlation with the RSA scores was more interpretable. The pattern of correlation change was moderated by lesion severity, revealing the sources of contribution to model performance. For the left lesions, left RSA unit correlations were much higher than the right RSA unit correlations in the milder lesion conditions. With increasingly severe lesions, the right RSA unit correlations increased with the decrease in the left RSA unit correlations. For the right severe condition, the left RSA unit correlations remained higher than the right RSA unit correlations. Fig. 4B shows that the intact model produced a higher RSA unit correlation for the LH1 than the RH1 and the opposite pattern was found for the left severe model. These results were conceptually similar to the findings of Fischer-Baum et al. (72), in which the healthy controls relied more on the left VWFA for orthographic processing in contrast to the patient with severe damage to left VWFA relied more on the right VWFA. Obviously, the tasks in the present study and Fischer-Baum et al.’s study are different. Thus, the patterns produced by the model can be considered as predictions for future patient studies in spoken production. Collectively, these results demonstrated that the RSA could potentially provide a more direct measure to relate model performance to the underlying computations.

**Table 2.**
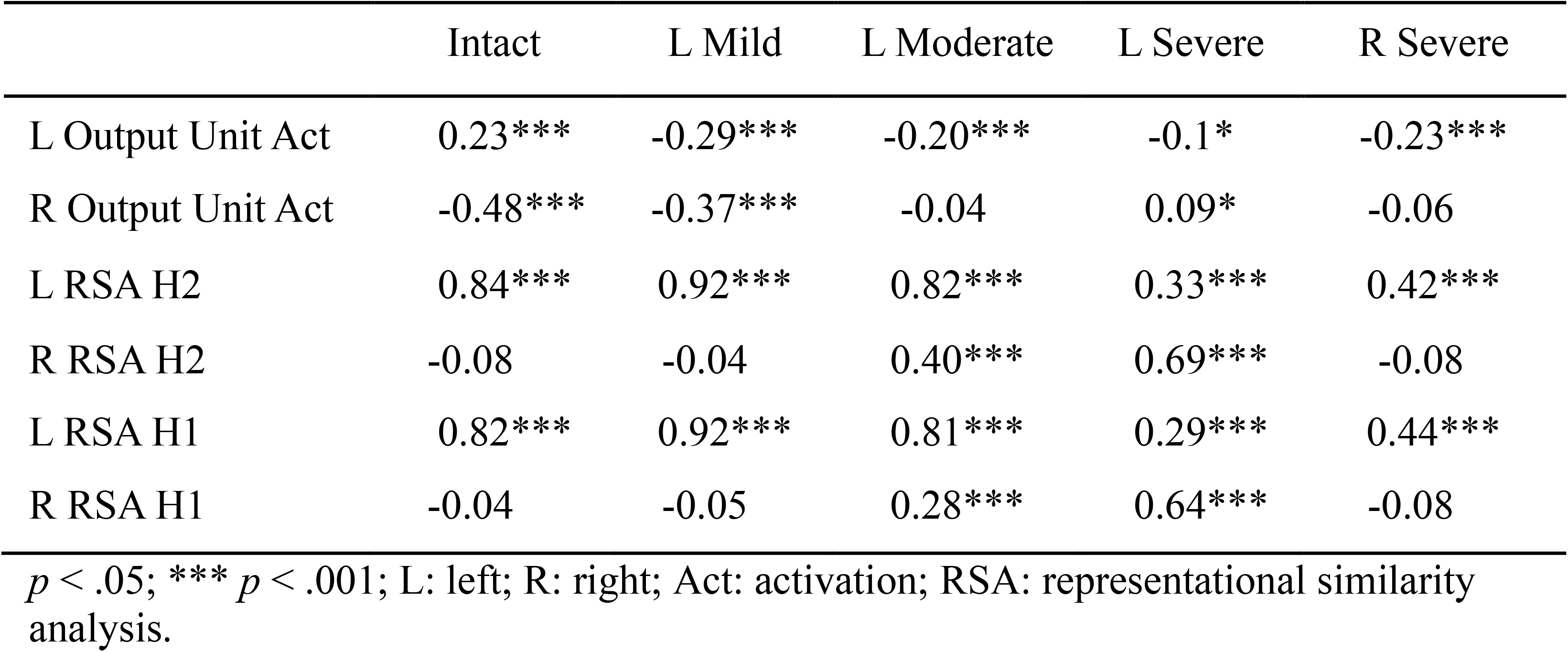
The correlations between model performance and output unit activations and RSA scores across the developmental learning period in the intact model and the re-learning period in the lesioned models

|  | Intact | L Mild | L Moderate | L Severe | R Severe |
| --- | --- | --- | --- | --- | --- |
| L Output Unit Act | 0.23*** | -0.29*** | -0.20*** | -0.1* | -0.23*** |
| R Output Unit Act | -0.48*** | -0.37*** | -0.04 | 0.09* | -0.06 |
| L RSA H2 | 0.84*** | 0.92*** | 0.82*** | 0.33*** | 0.42*** |
| R RSA H2 | -0.08 | -0.04 | 0.40*** | 0.69*** | -0.08 |
| L RSA H1 | 0.82*** | 0.92*** | 0.81*** | 0.29*** | 0.44*** |
| R RSA H1 | -0.04 | -0.05 | 0.28*** | 0.64*** | -0.08 |

$p < .05$ ; \*\*\* $p < .001$ ; L: left; R: right; Act: activation; RSA: representational similarity analysis.

### Interconnectivity between the left and right hemispheres

Thus far, the implemented model did not have interconnections between the left and right pathways. Cortical hemispheres, however, are connected by the corpus callosum as well as various subcortical routes (77). Given that the corpus callosum and interhemispheric connectivity are complex, a detailed neuroanatomically-constrained simulation is beyond the scope of this study. However, we explored a simplified simulation by adding direct ‘homotopic’ interconnections between the left and right pathways to investigate whether (a) this changed the patterns of simulated recovery reported above, and (b) if the model would develop transcallosal inhibitory connectivity as proposed in various classical hypotheses (6, 7, 9). Transcallosal connectivity in the model was implemented as sparse, bidirectional cross-connections between the left and right hemispheres without imposed positive or negative connections (all weight connections were allowed to develop freely). As there is no prior knowledge about inter-hemispheric connectivity density, we implemented two connectivity levels (30% and 70%). The training and testing procedures were exactly the same as previously described. We also ran additional comparison simulations in which the connections were constrained to be negative only *(see SI Appendix, S4)*. This constrained model produced stronger left lateralised patterns and it was less resilient to damage. Analyses of the weight values demonstrated that the vast majority were close to zero – i.e., the model effectively became one with independently functioning pathways. This was not true when the connection values were unconstrained.

Fig. 5 shows the resulting patterns produced by the left mild, left moderate and left severe and right severe lesioned models with different levels of interconnection (unconstrained). For comparison, the pattern produced by the model without interconnections is included in Fig. 5. Overall the patterns were similar. There was transient right unit activation for the left mild lesion condition but not for more severe left lesion conditions. In addition, the model could recover to a similar accuracy level regardless of the levels of interconnection. But, when the model had more interconnections, it showed a more bilateral pattern following damage and recovery, increasingly behaving like a single functional pathway model. This observation was confirmed by the results from the right severe lesion condition, where the model with more interconnections exhibited a more pronounced impairment in the early recovery phase.

**Fig. 5.**
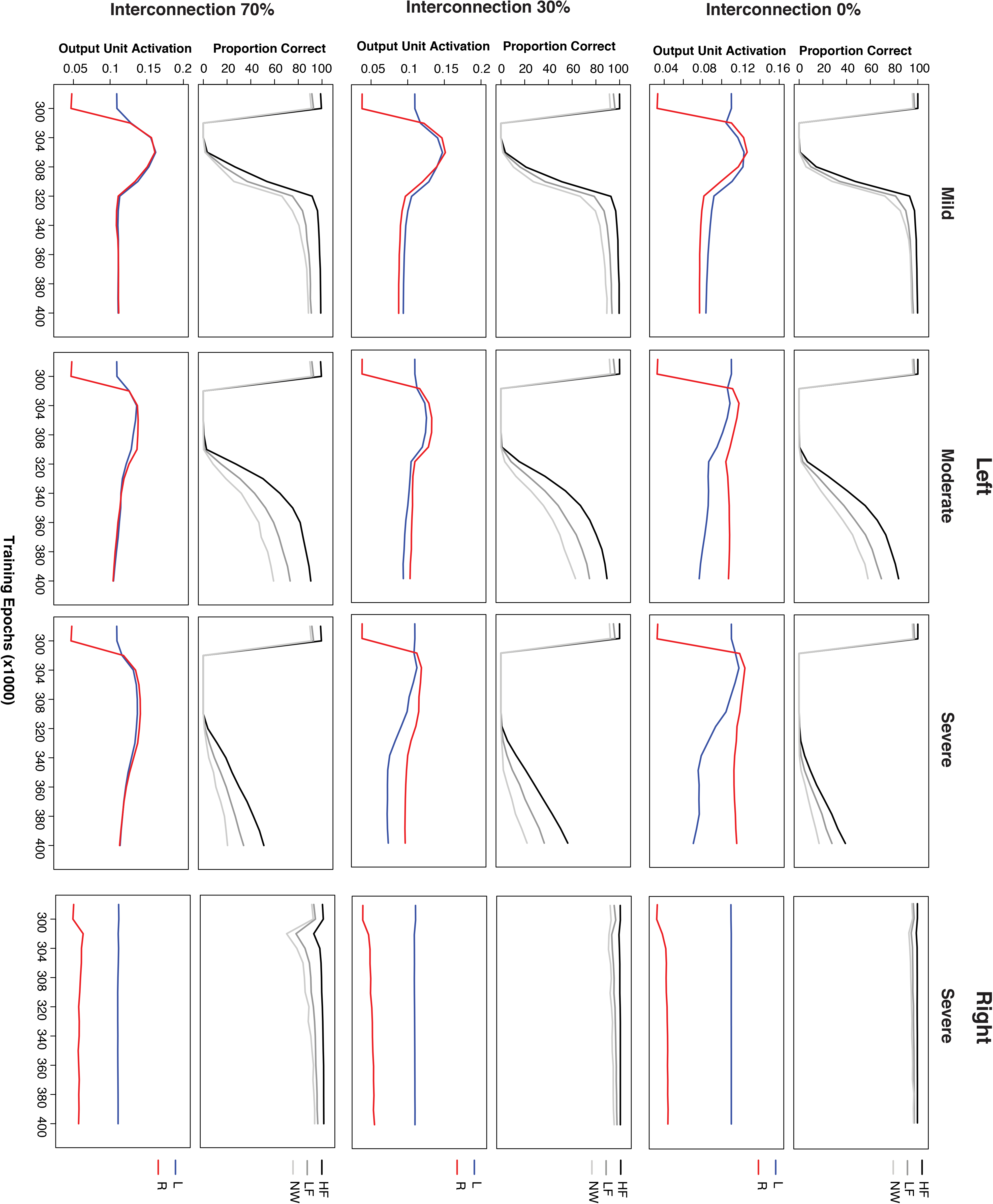
Simulation patterns of post-stroke aphasia and recovery produced by the model with three levels of interconnections (0%, 30% and 70%) between left and right sides for the left mild (20%[0.2]), left moderate (50%[0.5]), left severe (80%[0.8]) and right severe (80%[0.8]) lesion - conditions. The lesion level was a combination of the proportion (%) of the units was damaged and the range of noise (bracket) added to the connections to and from the hidden layer. For each lesion and interconnection conditions, the first panel shows model performance and the second panel shows output unit activation generated from the left and right pathway of the model separately. HF: high frequency words; LF: low frequency words; NW: nonwords; L: left; R: right.

## Discussion

Understanding the brain mechanisms underlying language processing is critical, both theoretically and clinically. To tackle various key issues that appear to be contradictory in healthy and impaired language processing (see Table 1), we developed a single, unified neurocomputational model of spoken language production with bilateral pathways. The key features of this modelling work include: the importance of considering healthy and impaired language within an intrinsically bilateral but asymmetric language network; to conceptualise recovery of function after damage as an experience-dependent plasticity-related learning process; and, to provide a platform to simulate behavioural and neuroimaging data from different populations. A list of the key findings is provided in Table 1 and each are discussed briefly below.

In an otherwise computationally-homogenous model, an initial imbalance in the processing capacity (number of hidden units) in the left and right hemisphere pathways was sufficient to explain the pattern of data observed in healthy participants and patients with chronic aphasia. Specifically, the capacity imbalance drives an emergent division of labour across the pathways such that the left hemisphere pathway picks up more of the computational work (i.e., each unit, on average, is more highly activated and contributes more to the final spoken output response than each corresponding right hemisphere unit). As a result, the undamaged model shows bilateral but asymmetric “activation” as observed in healthy participants. When this capacity imbalance is combined with plasticity-related recovery, the model provides an explanation for why left hemisphere stroke is more likely to result in chronic aphasia than right hemisphere stroke. Plasticity-related recovery reflects a re-optimisation of the remaining connection weights to maximise behavioural performance. This occurs in both ‘perilesional’ units and the contralateral pathway. The greater computational capacity in the left hemisphere means that, when the right hemisphere is damaged, there is greater capacity for the left hemisphere pathway to pick up the extra representational work previously undertaken by the (damaged) right hemisphere pathway (meaning that there is only transient aphasia). The same recovery process occurs following left hemisphere damage except that (a) the greater left hemisphere capacity means that, at least for mild levels of damage, there is still enough spare capacity in the remaining ipsilateral units to pick up the additional computational work (i.e., there is good or recovered function, and left hemisphere activation still dominates, even after mild levels of left hemisphere damage) and (b) there is insufficient capacity in the right hemisphere to compensate completely if the left hemisphere damage is too severe. In such circumstances, the model mimics chronic aphasia. In all cases, plasticity-related recovery means that there is a dynamic shift in the division of labour to ipsilateral ‘perilesional’ and contralateral areas, as is observed in fMRI studies of recovered patients. The model also demonstrates that there can be complex, nonlinear relationships between behavioural performance and levels of unit activation (a proxy for BOLD) whereas the relationship is much more direct when comparing performance to the accuracy of the representations coded in the pathway (implying that MVPA-type neuroimaging analyses may be a better way to assess and track the neural bases of recovery in aphasic patients).

Leftward hemispheric asymmetry has been shown in several brain regions and white matter tracts (36–40). However, there remains some controversy regarding a positive correlation between structural asymmetry and functional lateralisation (25, 41, 42, 78). The discrepancy could be related to individual differences among participants (e.g., age, education, handedness, and gender) or it could be because most studies have relatively small sample sizes (79). In a more controlled computational environment, our bilateral model with differential pathway revealed the impact of model capacity on the functional division of labour underlying performance and demonstrated a link between hemispheric asymmetry and language lateralisation. The simulation results are consistent with the positive correlation patterns reported in most right-handed healthy participants (41, 42). The model also shows that this structural difference could be fundamentally important for explaining patient data. By explicitly incorporating a leftward asymmetric but bilateral structure in the model, the model synthesises the seemingly contradictory patterns observed in both healthy participants and aphasic patients (Fig. 2): specifically, a leftward asymmetric but bilateral pattern in the intact model, and the much stronger lateralisation picture that is observed in chronic patients after left (aphasic) vs right (recovered) lesions. Additionally, the relationship between the severity of the left lesion and recovered performance is non-linear (see Fig. 2B), suggesting that the model had developed some resilience to mild damage (up to ~35%) but, beyond a “tipping point” the effects of damage cannot be overcome through plasticity-related re-learning, leading to more permanent language impairment as observed in chronic aphasia. It is important to note that there was a small divergence between the simulation results and Gajardo-Vidal et al.’s (17) patient data: that is, the model is more robust to right hemisphere damage (Fig. 2B). It is possible that this highlighted version of the model might possess a division of labour too biased to the left hemisphere pathway, making the contribution of the right hemisphere a little too weak. From our explorations of the key computational parameters, we know that the division of labour is governed in part by (a) the asymmetry in the balance of computational resources (Fig. 1) and (b) the degree of interconnectivity between the left and right pathways (Fig. 5).

Taking a step back, this explanatory framework for aphasia and recovery, raises some fascinating fundamental questions about higher cognition more generally: (a) why is it good for cognitive functions to be supported bilaterally; and (b) if so, why is it beneficial for some functions to remain at least partially dominated by one hemisphere (cf. the asymmetric yet bilateral architecture of the language production system)? Complete answers to these questions will have to wait for future research but there are some initial ideas in the literature. With regards to the benefits of bilateral implementation, a recent computational model and formal mathematical analysis demonstrated that bilateral systems are much more robust to the effects of damage than a singular system with the same resources (80). The second question is more difficult to answer. One possibility is that a bilateral system’s resources might be pulled asymmetrically if that cognitive function has to interact with other computations/representations that, themselves, are unilaterally expressed (though, of course, the begs the same question of why these are asymmetrically supported). A second possibility comes from a potential downside of distributing the same function across multiple brain regions. Whilst the distributed system may engender greater robustness to the effects of damage, it might induce a need for heightened synchronisation. Previous proposals (81) noted that human connected speech is highly demanding in terms of the rapid, accurate motor executions required, as well as the fast conceptual-to-speech transformations (82). When output signals need to change rapidly and accurately then, in the situation of bilateral systems, the need for synchronisation also increases (81). In the limit, sufficient synchronisation across distributed brain regions may be impossible to achieve and thus the ‘compromise’ is to let one side of the computation dominate, cf. an asymmetric, bilateral system.

Two potential mechanistic frameworks have been proposed for language recovery: degeneracy and variable neuro-displacement (13). Both mechanisms allow the language system to be at least partially resilient to damage and for recovery of function. Recovery can be accomplished by a permanent reformulation of the remaining multiple codes (degeneracy) or upregulation of systems (variable neuro-displacement), or both. The present neurocomputational model demonstrates that both mechanisms are not mutually exclusive and they can be utilised as a part of the recovery process. Immediately after dominant pathway damage, the model rapidly activates contralesional activation and also starts to re-formulate the perilesional unit contributions. If the perilesional units are capable of re-supporting the function, then later in recovery, both perilesional and contralateral contributions increase; otherwise, the perilesional contribution is decreased and the enhanced contralateral contribution continues. As such, it would appear that the recovery process follows the two proposed principles but the actual mechanisms involved depend on the level of task engagement by the units before damage and whether there is sufficient capacity in the remaining perilesional or contralateral areas to support recovery. As a result, there are differential output activation recovery profiles depending solely on lesion severity. With a mild left lesion, the perilesional units are largely persevered and can be re-formulated for recovery, leading to good recovery and left lateralised output activation patterns. With a more severe left lesion to the model, perilesional support is reduced and partial recovery relies mainly on the contralateral units. Accordingly, there is a co-occurrence of slow and imperfect recovered performance with right-lateralised activation patterns. These simulations collectively mirror the patient results reported in the literature: good performance is associated with left lateralised activations while worse performance is associated with more right lateralised activations (5); and, left-right-left changing brain activation patterns are observed in patients with mild brain lesions in the left hemisphere (51). This finding emphasises the importance of considering lesion severity when interpreting associations between good recovery and left lateralised brain activation patterns (5, 51) and the association between imperfect recovery and right-lateralised brain activation patterns (83).

The present bilateral model also provides a potential explanation for why the right hemisphere provides some but not perfect language support. The classical right hemisphere hypothesis (RHH) proposes that the right hemisphere is normally suppressed, via transcallosal inhibition, by the dominant left hemisphere system, but it can be released to provide some function after significant left hemisphere damage (6, 7, 9). As noted previously (2, 5, 8, 10–13, 27, 51, 57), the RHH leaves many puzzling questions open, including: how the RH can develop language representations under lifelong suppression; how left and right language systems might contribute to normal function; what bilateral yet asymmetric BOLD activation in healthy participants represents; and, why this biologically-expensive organisation for all people is an optimal solution for the minority of people who happen to suffer from the right kind of brain damage to induce aphasia. The current simulations provide a much more straightforward proposal for the data. The premorbidly bilateral albeit asymmetric system supports healthy function but can partially re-optimise following damage. This can all be achieved without any recourse to notions of juvenile RH language systems and interhemispheric inhibition. Instead, the RH subsystem is less efficient because it has less computational capacity and, in turn, learning in the left hemisphere over-shadows that in the right, resulting in the left hemisphere units taking up more of the representational work (Fig. 1C). These results follow even without interhemispheric connection. Even if included (Fig. 5), then (a) they do not all become inhibitory and (b) with increasing connectivity the model evolves into a single functional system. Of course, it should be acknowledged that the connections within corpus callosum are much more complex than the simple parallel connections implemented in the present model. Whilst interhemispheric connectivity has been shown to be inhibitory within the motor network (63–65), to our knowledge, there is currently no evidence of transcallosal inhibitory in language or other higher cognitive networks; in contrast, a few studies have demonstrated interhemispheric excitatory connectivity (66, 67). Finally, in a third variant in which inhibitory-only interhemispheric connections were enforced, the model set their value close to zero.

We should note that one previous study (9) applied TMS to left inferior frontal gyrus in healthy participants during a verbal fluency task, and showed decreased brain activity in the left but increased activity in the right homologue. These findings were interpreted as supportive evidence for transcallosal inhibition from the left to right hemispheres, however, the changes in the effective connectivity between the left and right inferior frontal gyri after TMS were not examined. Alternatively, the upregulation of homologue language areas after brain stimulation could be considered as a form of adaptive plasticity based on an interhemispheric compensatory mechanism (31, 32, 35, 84). For example, a recent study of semantic processing, combining theta-burst stimulation (cTBS) and DCM (32) found increased right ventral anterior temporal lobe (vATL) in response to cTBS to the left vATL. The DCM results revealed an increase in the facilitatory drive from the right to the left vATL. There was no evidence of negative inter-ATL connectivity with or without stimulation. Similar results have been reported in another brain stimulation study targeting Broca’s area during speech processing (35).

Lastly, the model investigated multiple measures within a single framework and their sometimes complex relationships. The simulation results suggest that, in task-based fMRI studies, BOLD signals and RSA measures may provide different information: although increase unit activations (cf. BOLD increases) are a necessary pre-cursor to behavioural recovery, higher unit activations do not necessarily imply that the units are contributing to improved performance. In the model, the performance improvement required both increasing unit activation and tuning weight connections. Immediately after damage, the activation level of the units in the model was generally low. Thus, the first step toward re-learning was to increase the activation level via a generalised weight connection increase. This was followed by re-tuning weight connections in order to minimise the errors between the target and actual patterns at the output layer. The implication is that fMRI BOLD signals in patients during recovery have an ambiguous interpretation; they could reflect the neural basis for recovered performance (as occurs, for example, in the right hemisphere pathway after severe left hemisphere lesions: BOLD-type and then RSA-type measures increase, see Fig. 3) or alternatively generalised but untuned activation (e.g., after a moderate lesion: right hemisphere activation increases but performance recovers only after the left hemisphere RSA has bounced back, see Fig. 3). In contrast, RSA might provide a more direct measure to link recovered performance with neuronal pattern information in different phases of aphasia recovery. This result is consistent with a growing interest in using different types of neuroimaging analyses to investigate the right hemisphere activation patterns in post-stroke aphasia and how it is related to recovered performance (71, 72). Given that, to date, there have not been any RSA-based studies of spoken language production in aphasic patients, the current simulations serve as model predictions for future neuroimaging studies. By extension, the same techniques might also be helpful in clarifying the (dis)advantages of using brain stimulation techniques (TMS or tDCS) to alter brain activation for effective treatments.

To finish we note explicitly that there were at least three deliberate simplifications adopted in the model which can be addressed in future work. First, the model focused on speech production along the dorsal pathway. Obviously, there are multiple pathways in the language network (16, 40, 85–88). For example, we have not considered the ventral pathway that includes a semantic system for comprehension. A previous neurocomputational model (46) demonstrated that a dual-pathway neural network model could simulate different types of aphasia (including receptive and expressive language) based on damage to a corresponding lesion site. Secondly, the psycholinguistic and cognitive science literatures contain many sophisticated models of individual language tasks (typically in relation to healthy performance) that embrace large word corpora and linguistic detail. We deliberately adopted a much more psycholinguistically-simple model so that we could more easily explore the potential relationship between behavioural performance and brain structures/pathways, and thus allow us to distil some principal mechanisms about the emergent division of labour between the two hemispheres and how this can change in recovery. Third, we acknowledge that the backpropagation algorithm has commonly been viewed as biologically problematic even though it is the most efficient and effective for deep neural networks (89). However, this view may be changing: a very recent review (90) proposed that there could be a more direct neural analogue of backpropagation: in simple terms, top-down feedback/prediction could be compared at the local level against the bottom-up sensory input. Clearly, however, more empirical evidence connecting the backpropagation algorithm and learning in the brain is needed. Future models can merge and elaborate these approaches to provide further systematic investigations, thereby elucidating the neural bases of healthy language and partial recovery in post-stroke aphasia.

## Methods

### Model architecture

The bilateral model was implemented as a simple recurrent network. The dual pathway architecture of the model is shown in Fig. 1A. Each processing pathway consisted of two hidden layers. For the first hidden layers, there was a copy of the hidden layer, shown as dashed lines in Fig. 1A, known as an Elman layer (91); this allows a copy of the hidden layer from the previous time tick to influence the current unit activations, functioning as a memory buffer in the model.

The input phonological layer was connected to the first left and right hidden layers with Elman connections and then to the second left and right hidden layers and then to the single, final output layer. All layers were fully connected and the connections were unidirectional unless stated otherwise.

### Representation

100 three-phoneme high frequency and 100 three-phoneme low frequency monosyllabic words with consonant-vowel-consonant structures were included in the training set. Each word was represented by three phoneme slots, with each slot consisting of 25 phonetic features (92, 93) *(for details, see SI Appendix, Methods).* The nonword list comprised 25 items creating by changing the first consonant, the vowel or the final consonant in a word.

### Training and testing

The model was trained on word repetition. For each word, the model was run for six time ticks. In the first three time ticks, each phoneme was presented in the input layer sequentially. There was no output target until all the phonemes were presented. From the fourth time tick to the sixth time tick, the model was required to produce the target phonemes sequentially. Which word was presented to the model was determined by its logarithmic frequency (94). The model was trained using a standard back-propagation algorithm. Weight connections were updated after each word presentation on the basis of cross-entropy error.

All words were used for training and the nonword list was used for testing generalisation. The model was trained for 300,000 word presentations, at which point the model repeated words accurately (> 98%) and generalised to nonwords (> 96%) (Fig. 2A). The model’s performance was assessed by comparing the output phonological pattern with the sequential phoneme target. The model was judged to be correct only when all phonemes were correct. Twenty versions of the model with different random initial weights were trained to simulate different participants and to prevent the results emerging from idiosyncratic random initial weights. More detailed training environment and testing procedures are reported in *SI Appendix, Methods*.

### Neuroimaging correlates

In neuroimaging studies, a laterality index was computed by subtracting the signal obtained in the right language areas from the corresponding left language areas and then dividing the score by the sum of the signals (42, 74). For the simulation, two measures were used: one was functional correlation (47, 75) as a proxy of effective connectivity signals, and the other was output unit activation (76), as a proxy of fMRI activation. For functional correlation, we recorded the unique contribution from the left pathway to the output phonological layer for all words (from the fourth to the sixth time ticks). This was achieved by lesioning the links between input and the right hidden layer 1 so that there was no input signal from the right pathway. Similarly, the right pathway contribution was obtained by lesioning the links between input and the left hidden layer 1. We then correlated the activation patterns from each pathway with the patterns when both pathways were utilised. For output unit activation, the same lesioning technique was used to isolate the unique contributions from the left or right pathway to the output layer activations. Both measures were used to compute the lateralisation index (a positive score indicates a left-lateralised pattern).

### Representational similarity analysis

For each word, the model produced three phonemes sequentially. To conduct representational similarity analyses (95), we concatenated the three output activations into one output pattern for each word. We then computed a target representational dissimilarity matrix (RDM) based on the correlation distance of all word pairs. Similarly, for model RDMs, we computed the matrices based on the correlation distance of hidden unit activation patterns at hidden layers 1 and 2 in the left and right pathways of the model independently. The hidden unit activation pattern for each word consisted of hidden unit activations of the constituent phonemes. The RSA correlation scores between the target RDM and the model RDMs of hidden unit activations were reported.

## Acknowledgements

This research was supported by an ERC Advanced Grant to MALR (GAP: 670428 - BRAIN2MIND_NEUROCOMP) and Medical Research Council intramural funding (MC_UU_00005/18).

## Code availability

The computer scripts used to run the simulations and to analyse the results are available on the MRC Cognition and Brain Science Units Data Repository (https://www.mrc-cbu.cam.ac.uk/bibliography/opendata/)

## Data availability

The datasets generated and analysed in this study are available on the MRC Cognition and Brain Science Units Data Repository (https://www.mrc-cbu.cam.ac.uk/bibliography/opendata/)

